# SEEDS: Simulating Emergence of Errors in DNA Storage

**DOI:** 10.1101/2025.02.14.638249

**Authors:** Mst. Fahmida Sultana Naznin

## Abstract

**Background:** DNA storage is a nonvolatile memory technology for storing data as synthetic DNA strings which offers unprecedented storage density and durability. Yet, the application of DNA as a practical digital information storage medium remains an enigma, since this is extremely expensive and it takes a substantial amount of time to encode and decode data to/from synthetic DNA. More importantly, various phases of DNA storage pipeline (e.g., synthesis, sequencing, etc.) are error prone. Furthermore, DNA is subject to decay over time and the reliability of the synthetic DNA depends on various aspects, including preservation medium and temperature. To allow for the perfect storage and recovery of the information and thereby making it competitive with the existing flash or tape based technologies, advanced error protection schemes are necessary. However, evaluating and comparing various DNA storage technologies and error correcting codes under realistic model conditions – comprising a wide array of synthesis medium, sequencing technologies, temperature and duration – is prohibitively time consuming and expensive.

**Results:** In this study, we present SEEDS, an error model based simulator to mimic the process of accumulating errors at different phases of DNA storage. SEEDS is the first known simulator which incorporates various empirically derived statistical (or stochastic?) error models, mimicking the generation and propagation of different types of errors at various phases in DNA storage. It was assessed for its validity against the data from a number of published wet-lab experiments.

**Conclusions:** SEEDS is easy to use and offers flexible and comprehensive parameter settings to mimic the error models in DNA storage. Validation against in vitro experimental results suggests its promise for emulating the stochastic models of error generation and propagation in DNA storage. SEEDS is available as a web interface with a server side application, along with portable cross-platform native applications (available at givethelink).

## 1 INTRODUCTION

We are living in an age of information, where the world is producing quintillion (∼ 10^18^) bytes of data each day [64]. To handle this ever-increasing amount of data, we need dependable storage technologies because traditional magnetic and optical storage devices are not reliable for long-term data storage (> 50 years) [85]. Besides, hard disks are expensive and they require a constant supply of electricity. Moreover, building and maintaining data centers to store data on tape drives would require lots of efforts, power, time, and money [29]. This is an ever-growing problem, particularly in the field of life sciences, where petabytes (∼ 10^15^) of digital information are being produced every year by genome sequencing alone [20, 32]. Therefore, it is time to look beyond the traditional storage devices. Luckily, a prospective candidate has emerged from the field of life sciences itself, in the form of *DNA storage*.

DNA is Nature’s own medium to store hereditary information and genetic instructions across species. DNA is attractive as a storage medium for its capacity to store gigantic amount of data within its negligible space. Theoretically, DNA can encode at most two bits per nucleotide or 455 exabytes per gram of ssDNA [18]. Despite being highly dense, DNA storage requires very low maintenance power (< 10^−10^ watts per gigabyte) [29] and have greater longevity (hundreds of thousands of years) than any other available storage media [6, 19]. Therefore, DNA offers an attractive target for information storage.

At first, in 1988 a 35 bit binary sequence of a graphic image was encoded into bacterial DNA [22]. After that, over the years many attempts have been taken to improve the scalability and performance of DNA storage [13, 15, 18, 37, 105]. One of the greatest breakthroughs came when researchers from Microsoft and University of Washington were able to achieve random access for large-scale (200 MB) data storage [74]. Even that was surpassed when Erlich and Zielinski were able to achieve enormous data density, storing around 215 petabytes of data per gram of DNA, which they termed as DNA fountain [25]. Even recently researchers from North Carolina State University have developed new techniques for selectively extracting unique files from complex database of DNA by mimicking as much as 5 terabytes of data [96].

Despite all these advantages and progresses, DNA storage has its own shortcomings as to err is any storage device. To store data in DNA and then retrieve it later, the data need to go through several phases. Firstly, the binary information is encoded into nucleotide sequence. Then the sequences are synthesized into DNA (writing data) and stored into suitable medium for long term usage (Figure 1). Finally, when one needs to retrieve the data, sequencers are used to produce reads from the stored DNA fragments (reading data). The nucleotide sequences generated from the reads are then decoded back to binary information.

**Figure 1.**
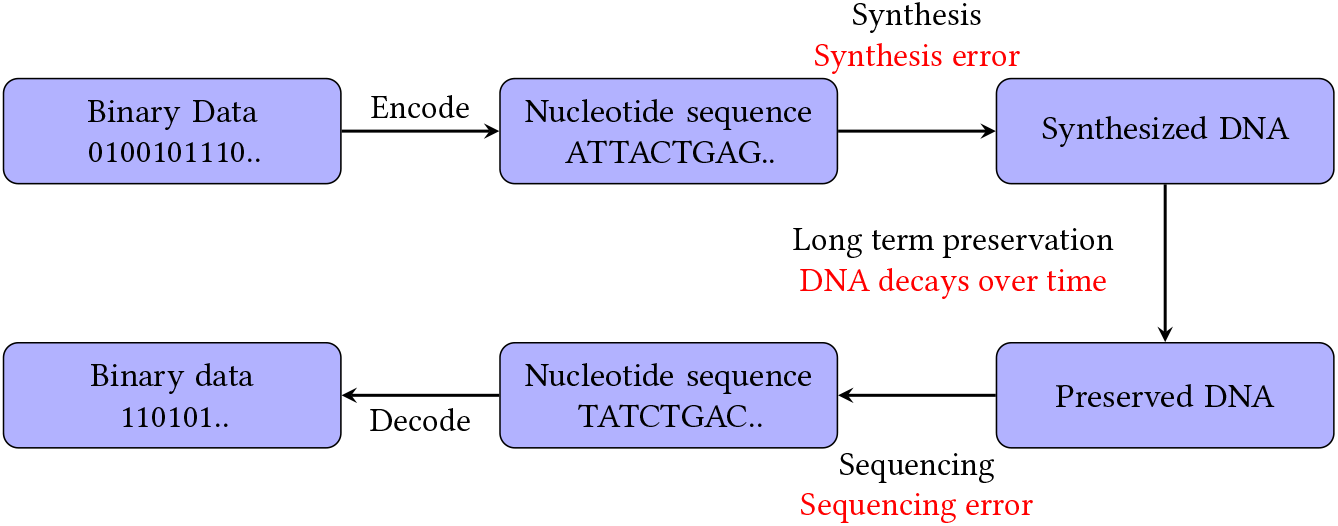
Different stages of storing information via DNA storage. Bayzid: I would prefer a more fancy figure. We will discuss this in our next meeting.

As data traverse all these steps in the form of DNA, errors accumulate in each phase. Both DNA synthesis (writing) and sequencing (reading) are far from perfect, with error rates as often as about 1% per nucleotide [15, 29]. Besides, DNA goes through degradation over time [4]. All these make DNA more vulnerable to errors than traditional storage devices. Therefore, to realize an efficient DNA storage mechanism, many studies have been conducted particularly aiming at developing robust error-correction mechanisms for DNA storage [13, 38, 104]. As DNA might be the future for next generation data storage, further research is required to explore and elicit its potential to the full.

However, the efforts in this regard are often capped by the costs associated with DNA storage. The relatively high cost of DNA synthesis (writing) makes it problematic if DNA storage is going to be the standard archiving technology. Back in the 1980s, it took $6, 000 to synthesize only 10 nucleotides [72]. However, over the next few decades oligonucleotide synthesis became more routine and with the advent of new technologies, the cost came down to less than $1 per base [27, 33]. Yet if we consider the cost of synthesis as low as 7 cents per bp (offered by Twist Bioscience [12]), it would still cost us around $100, 000 to store a single minute of high quality stereo audio [67].

On the contrary, the cost of DNA sequencing (reading) has reduced over time [47]. Even the feat of sequencing human genome has hit the mark of $1, 000 [56]. Still the cost is too high for application level implementation considering the cost of instrument, cost per run, and reading cost per unit of data. Though there are portable, cheaper sequencing technologies available, e.g., Oxford Nanopore, error rates are too high among the produced reads to be viable [50, 97].

Besides writing and reading DNAs, we need to preserve them within suitable environment so that the stored information can be retrieved properly without much damage. It has been observed that high temperature works as a factor for DNA degradation [52]. Therefore, to maintain the integrity of DNA over long periods of time, we need storage conditions (e.g., low temperature or dry state) in which molecular mobility is more restricted. However, maintaining samples at low temperatures is exceedingly costly requiring hundreds of millions of dollars to purchase the necessary freezers and maintain the frozen samples [24]. On the contrary, though storing dehydrated or solid-state DNA at room temperature can significantly reduce preservation cost [6], stored DNAs go through degradation within a few years [49].

Given all these costs at hand, DNA storage is not cheaper at all. Earlier in 2013, Nick Goldman and his team encoded 739 KB of information into DNA. They estimated a cost of $12, 400 MB^−1^ accompanied by decoding cost of $220 MB^−1^ [37]. On the other hand, the chemical preservation technique proposed by Robert N. Grass and his team to store digital information in silica-encapsulated DNAs with error correcting codes cost as much as about $2, 500 for synthesizing DNAs to store 83 KB of information [38]. Erlich and Zielinski’s DNA fountain project cost around $10, 000 to store 2.15 MB of information [25]. Moreover, the portable, error-free DNA storage mechanism proposed by Yazdi, Gabrys, and Milenkovic cost $2, 540, apart from sequencing cost, only to write (synthesize) two images into DNA (3.6 MB of data) [104]. Such high costs often affect the flow and outcome of research activities particularly for countries with limited resources and research facilities [5].

Therefore, it is evident that the usefulness and performance of DNA storage are shaped by the costs as well as errors associated with synthesis, sequencing, and preservation of DNA. Now, the reduction in both costs and errors is dependent on the advancement of technology. Therefore, to facilitate various research activities, particularly aimed at finding effective encoding, decoding, error-detection, and error-correction mechanisms for DNA storage, this work introduces the idea of a simulator that would model the errors that occur in different phases of DNA storage.

Recently many computational tools have been developed for simulating next-generation and third-generation sequencing processes [3, 26, 68, 91]. Besides, a couple of simulators have been designed to simulate the encoding and decoding phases of DNA storage [2, 15, 86, 93]. These simulators take a text file as input and encode them into DNA string, i.e., nucleotide sequences and again decode them back to original string. Some of them evaluate the performance and reliability of different encoding mechanisms [15], whereas, some others report various biochemical properties of encoded DNA sequence, such as GC content, melting temperature, and total cost to store files in DNA [86]. Others simulate how digital information can be encoded and stored into various bacterial systems [2, 93].

However, to the best of our knowledge, there exists no such computational tool that captures how errors propagate during synthesis as well as preservation phases of DNA storage. In this study, we present SEEDS (Simulating Emergence of Errors in DNA Storage) to facilitate modeling of different types of errors at various phases in the pipeline of DNA storage. To the best of our knowledge, SEEDS is the first work that attempts to combine various comprehensive mathematical models of errors at different phases of DNA storage into a single pipeline.

SEEDS will provide an economic and flexible way to evaluate different hypothesis, encoding-decoding mechanisms, error-detection, and error-correction techniques related to DNA storage prior to wet-lab implementations, which are usually costly and time consuming. We believe that such a tool would help the researchers to develop a clear understanding about how errors accumulate in DNA storage, and to use this knowledge in designing new methodologies. The integration of various types of errors through a stochastic mathematical model would enable us to evaluate the performance of various synthesis, sequencing, and preservation techniques as well as to choose optimal hyper-parameters/conditions for effective performance of DNA storage.

## 2 METHODS AND IMPLEMENTATION

SEEDS was designed in Java to facilitate the features of object-oriented programming to ensure effective, portable, and robust simulation techniques across all platforms. It is a platform-independent software designed to run on standard desktop computers/ laptops with integrated Java Runtime Environment (JRE). The simulator uses a graphical user interface, based on Swing, to interact with the user. It enables the user to give input an encoded nucleotide sequence, then works on that sequence based on different errors that are likely to take place at different stages of DNA storage. We implemented various stochastic models of errors as separate independent modules, such as, synthesis module, preservation module, and sequencing module. The selection of different stochastic models of errors in DNA, across different experimental settings, was shaped by the availability of such data. These modules take input a string of nucleotide sequences and introduce different types of errors corresponding to that stage of DNA storage. The basic skeleton of our simulator is as follows:

- **Input:** a string of nucleotide sequences
- Pass the input nucleotide sequence through:
  – Synthesis module()
  – Preservation module()
  – Sequencing module()

- **Output:** The system generates following outputs:
  – Error rates for each nucleotide
  – A log file indicating the type of error *T* that affected a nucleotide *N* at base pair *X*
  – Half-life of DNA, average fragment length, bond survival rate during preservation
  – Modified nucleotide sequence with errors
  – Graphs depicting error rates, average fragment length, bond survival rates, etc.

An input DNA sequence goes through different modules corresponding to different phases of DNA storage over multiple iterations as selected by the user. For example, *synthesis module* corresponds to the synthesis phase of oligonucleotides and introduces necessary errors in the input nucleotide sequence that take place during DNA synthesis. Next, the string is passed through *preservation module* where errors and random fragmentation are introduced depending on the length of preservation period and temperature. In the end, the modified nucleotide sequence reaches *sequencing module* where errors that are likely to occur during reading (i.e., sequencing) are integrated. The errors that accumulate in these phases can be divided into 4 main categories:

1. **Deletion**: In this type of error, a part of DNA sequence is lost either during synthesis, sequencing, or preservation phase of DNA.

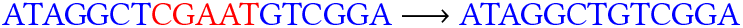
2. **Insertion**: This is the addition of one or more nucleotide base pairs within a DNA sequence.

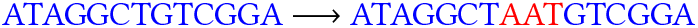
3. **Transition**: Transitions are interchanges of two ring purines (A ↔G) or one-ring pyrimidines (T ↔C).

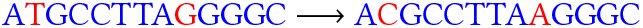
4. **Transversion**: Transversions involve interchanges of purines (A, G) with pyrmidines (T, C).

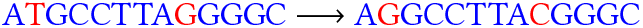

In general, we integrated many of these errors using probabilistic models based on experimental data and and various String manipulation techniques. For errors, such as transition and transversion, we replaced a nucleotide base with another as applicable and determined by the weighted probabilistic distribution of the nucleotide bases for these errors. We also included some other subtle modeling capabilities, such as, introduction of custom error rates based on extrapolation of existing experimental error rates, implementation of biases in errors towards certain ends of the genomic sequences, modeling metabolic nature of DNA at varying temperatures, and so on. Finally, the modified nucleotide sequences/ fragments containing errors are listed in an output file named ‘output.txt’. We also generated a log file (‘log.txt’) that contains bp positions and nucleotides associated with errors in different phases of DNA storage. Besides, we created a text file named ‘results.txt’, which contains error rates of different types of errors and other measures (i.e., half life of DNA, average fragment length, bond survival rate, etc.,) related to different phases of DNA storage. The details of different error modules of our simulator are discussed in detail as follows.

### 2.1 Synthesis Module

With the rise of synthetic biology, we have embarked on an era of creating new functional genes by enzymatic assembly of chemically synthesized oligonucleotides [21, 95]. Bayzid: the following line is hard to follow: Mahjabin: Sir, please check if it alright The resulting genes often contain errors due to flawed chemical oligonucleotide synthesis and faulty enzymatic gene assembly processes. Current gene synthesis technologies have typical error rates in the range of 10^−2^ to 10^−3^, or 1 − 10 errors per kilo base pairs synthesized [17, 45, 94, 103]. For our synthesis module, we incorporated error rates from a number of available synthesis technologies. The basic steps followed by our synthesis module is as follows:

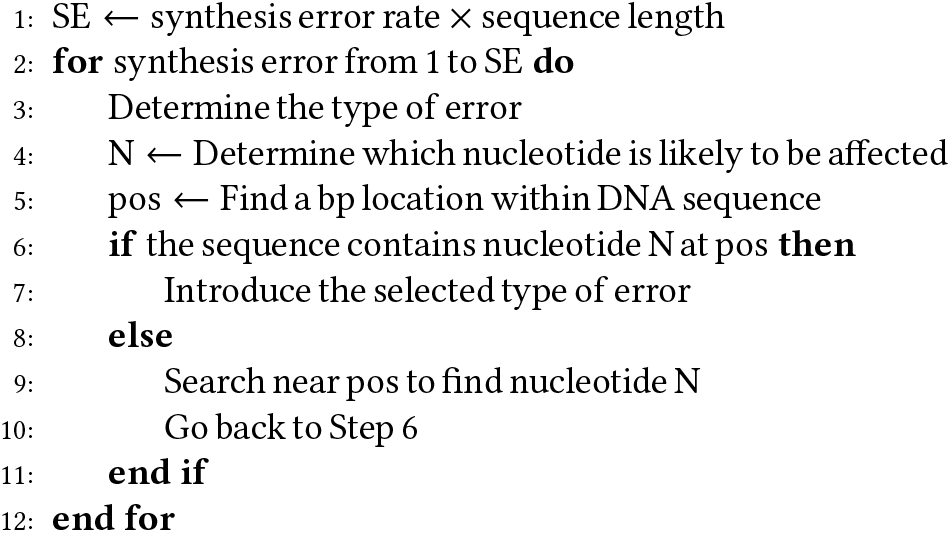

In the following, we explain these steps for different gene synthesis technologies as incorporated into our synthesis module.

#### (1) *De* novo DNA synthesis

This technology is based on a combination of organic chemistry and molecular biology techniques, where complete gene sequences may be synthesized *de novo* without any template DNA. Over the last half century, *de novo* DNA synthesis has seen significant improvements in terms of throughput [34, 35]. However, given an error rate, the probability of finding an error-free synthetic DNA sequence decreases exponentially as its length increases [62]. Error rates for *de novo* technology vary in the range of 1 in 600 bp (equivalent to 1.67 errors in 1000 bp) to 7 errors in 2.2 kbp [10]. Typical error rates fall in the range of 1.5 to 1.8 errors per 1000 bp [17, 45, 102]. Due to such varying error rates from experiment to experiment, we used a normal distribution (mean error rate 1.66 per 1000 bp, SD = 0.00015) to produce error rates for ‘*de novo* DNA synthesis’ module of our simulator. We obtained this distribution from the error rates reported in previous experiments. We used this error rate along with sequence length to calculate the number of possible synthesis errors in step one. Next, we used a stochastic model to introduce all these errors into the given DNA sequence (step two).

In this regard, our first challenge was to identify whether a particular synthesis error would be either deletion, insertion, transition, or transversion (step three). For this purpose, we used empirical distribution of different types of errors (Figure 2a) based on experimental outcomes from the work of Carr *et al*. [17]. From Figure 2a, we can seen that, in the wet-lab implementation of *de novo* DNA synthesis without any error correction, deletion errors are more abundant (65%) than other types of errors. On the other hand, insertion, transition, and transversion errors contribute to 5%, 16%, and 14% of total errors respectively. We used this error distribution to determine the type of a particular synthesis error.

**Figure 2.**
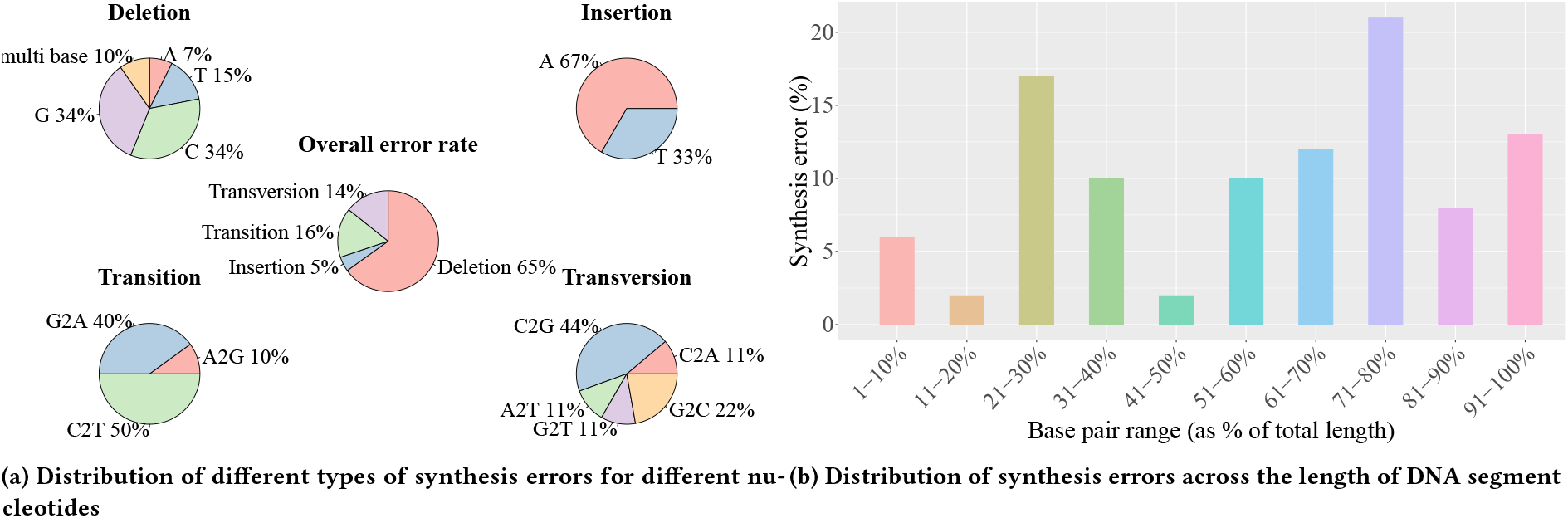
Empirical error rates for *de novo* DNA synthesis without error correction mechanism from the work of Carr *et al*. [17]. Figure (a) represents distribution of different types of errors (insertion, deletion, transition, and transversion) for different nucleotides (A, T, C, G) along with overall distribution of the errors. Figure (b) represents distribution of synthesis errors across the length of synthesized DNA sequence. This figure shows percentage of synthesis errors that fall within a particular base pair range of the synthesized DNA sequence.

**Figure 3.**
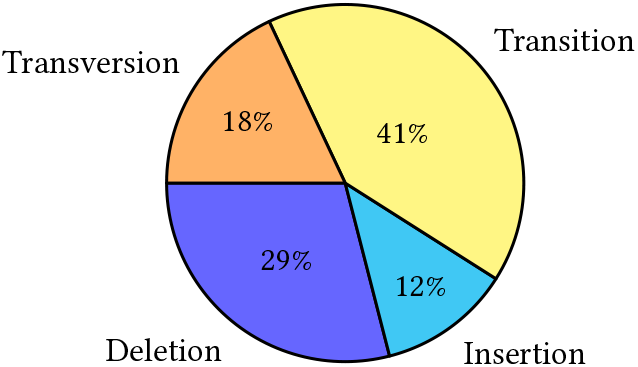
Empirical error rates for microchip based DNA synthesis technology from the work of Tian *et al*. [94].

Now, since there are four nucleotide bases, the selected type of error may involve any one of them. Therefore, we further looked into the distribution of nucleotide bases across different errors from the work of Carr *et al*. [17]. Firstly, in case of deletion (Figure 2a), cytosine and guanine are more likely (34%) to be deleted, whereas, adenine (7%) and thymine (15%) have moderate error rates. Besides, according to their study, 10% of the deletion errors involve multiple bases. Secondly, in their experiment, 50% of the transition errors involved transition from cytosine to thymine. Transition between purine bases contributed the remaining (G →A: 40%, A →G: 10%). Thirdly, among different types of transversion errors, transversion between cytosine and guanine are more common (C →G: 44%, G →C: 22%). Transversions from adenine to thymine (11%), guanine to thymine (11%), and cytosine to adenine (11%) are also observed (Figure 2a). Finally, in case of insertion, only adenine (67%) and thymine (33%) were involved in *de novo* DNA sequences synthesised by Carr *et al*. [17]. Therefore, after determining the type of error, we used weighted probabilistic distribution of different nucleotide bases to figure out which nucleotide base is more likely to be affected by that particular type of synthesis error (step four).

Next, we needed a particular location within the DNA sequence containing the particular nucleotide to introduce the selected type of synthesis error. Following the experimental outcome of Carr *et al*. [17], we observed that synthesis errors are not equally distributed across the length of the synthesized DNA sequences. Some segments of DNA contain more errors than others. The empirical distribution of observed errors across the length of synthesized *de novo* DNA sequences in [17] is shown in Figure 2b. It can be observed that majority of the synthesis errors (21%) take place in a base pair (bp) region that fall within 71− 80% of the original DNA sequence. Base pairs that fall within 11− 20% and 41− 50% of the synthesized DNA sequence, encounter the least (2%) amount of synthesis errors. Around 64% of all synthesis errors take place within the last 50% of the DNA sequence, whereas, the base pairs at the beginning have less errors. We used a weighted probabilistic distribution of errors within different segments of DNA to estimate particular bp location to introduce the selected type of synthesis error (step five).

However, the selected bp location might not contain the nucleotide that we selected to introduce synthesis error (step eight). In this case, we search at both ends of the selected bp location to find the desired nucleotide base (step nine). Finally, when we found the desired nucleotide base either at or near the selected bp location, we modified the DNA sequence following the selected type of synthesis error (step six and seven).

#### (2) Microchip based DNA synthesis

Photochemical reactions dependent [11, 107] microchip based gene synthesis technology has seen significant improvements in terms of cost, efficiency, and throughput over the past couple of years. Despite low cost, oligonucleotides synthesized on microarrays are prone to higher error rates [63]. Therefore, different techniques have been adopted to reduce the error rates of synthesized oligonucleotides [81]. We utilized error rates from previous works on microchip based DNA synthesis for this part of the synthesis module of our simulator.

The error rates vary in the range of 1 in 1394 bp [94] to 1 in 1500 bp, 1 in 1130 bp, 1 in 1350 bp [55], and so on. We used a normal distribution (mean error = 1 in 1329 bp, SD = 0.0000936) of these reported error rates to generate error rate for this part of the synthesis module of our simulator. We used the error rate generated from this distribution with sequence length to calculate the number of possible errors in step one.

Next, to introduce each of these errors (step two) we required to know the distribution of different kinds of synthesis errors in microchip based DNA synthesis technology. In this regard, we used the empirical distribution of various errors (3) reported in the work of Tian *et al*. [94]. We particularly chose this work for the detailed mention of different kinds of errors. From 3, it can be seen that, among all the errors, transition error is the most abundant (41%), while deletion (29%) and transversion (18%) errors are present moderately. Furthermore, insertion errors again contributed the least (12%) of all synthesis errors. We used weighted distribution of these errors to randomly select the type of synthesis error (step three).

However, we did not come across any relevant work with the distribution of different nucleotide bases for different types of errors in microchip based DNA synthesis. Hence, we randomly selected a nucleotide in step four, assuming that each of them is equally likely to be affected by the error. Moreover, we could not find any work that reported the frequency of synthesis errors across the length of synthesized oligonucleotides. Therefore, in step five, we randomly selected a bp location within the DNA sequence assuming that the synthesis error is equally likely to be present at any part of the DNA. If the selected location contained the desired nucleotide (step six), we modified the sequence following the type of selected error (step seven). Otherwise, the neighbourhood of the selected bp location was searched until the desired nucleotide was found(step nine, ten).

#### (3) Protein-mediated error correction for *de novo* DNA synthesis

This method deals with error reduction for *de novo* gene synthesis. This approach is based on a protein that is part of DNA mismatch repair pathway. This protein binds to many different kinds of DNA mismatches [66] as well as short deletions or insertions of one of four nucleotide bases [101]. Peter A. Carr and his team manipulated this affinity for mismatches to reduce errors from the desired products in *de novo* DNA synthesis [17]. Using this approach, they have been able to achieve an error rate of 1 per 10000 bp, which is 15 fold lower than typical error rates in *de novo* DNA synthesis. Our work uses this error rate to calculate the number of probable synthesis errors in step one. Secondly, to incorporate each of these errors (step two), we used a random probabilistic model based on the empirical distribution of errors (Figure 4a) in this method to determine which type of error is more likely to take place (step three). As we have already mentioned, this method produces very little error than traditional *de novo* DNA synthesis. In their wet-lab implementation of this error correction method, they sequenced a total of 39080 bps which contained only 4 synthesis errors. Among these 4 errors, two (50%) were transversions (G →C, C→ A), one (25%) was a deletion of adenine, and another one (25%) was a transition from guanine to adenine. These distributions were used with a probabilistic model to figure out which nucleotide is likely to be affected by the error (step four). Next, a bp location was selected within the DNA (step five) based on the empirical distribution of errors across different bp range within the sequence (Figure 4(b)). Among the 4 synthesis errors in their experiment, two fell within first 1− 10% of the bps, whereas, the remaining two were within a bp range that fell within 21 − 30% and 61 − 70% of the bp regions respectively.

**Figure 4.**
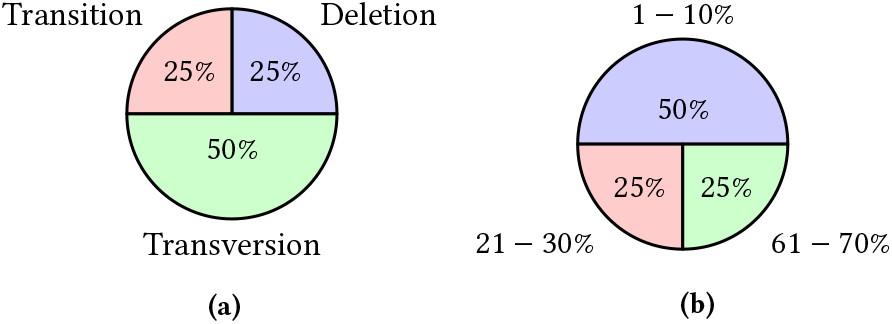
Empirical distribution of errors using protein-mediated error correction for *de novo* DNA synthesis from the work of Carr *et al*. [17]. Figure (a) represents distribution of different types of errors. Figure (b) represents distribution of these errors within different bp range of the synthesized DNA sequence.

Afterwards, if the selected bp location contained the desired nucleotide, the given DNA sequence was modified in accordance with the selected type of error (step six, seven). Otherwise, the neighbourhood of the selected bp location was searched until one with the desired nucleotide was found(step nine, ten).

Bayzid: Place this para under the heading “custom error rate”. Alternatively, move the first two sentences from “custom error rate” out to this para. Mahjabin: Sir, it is done

#### (4) Custom error rate

Besides incorporating errors from existing gene synthesis technologies, options were provided so that the users can use varying error rates that suit their purpose. Synthesis errors vary a lot from experiment to experiment depending on DNA synthesis technologies and error correction mechanisms. Therefore, this option was introduced to accommodate any future change in the error rates of synthesized oligonucleotides and also to enable users to customize the performance of the simulator to suit their own needs. This option enables users to give input varied synthesis error rates (per bp). This error rate is then directly used in step one to calculate number of possible synthesis errors. Next, simple linear models were used to determine the type of each of these errors (step three). These models were developed based on the distribution of different kinds of errors in *de novo* DNA synthesis [17] (Figure 2a), microchip based DNA synthesis [94] (3), and protein-mediated error correction for *de novo* DNA synthesis [17] (Figure 4a) as described above. Given an overall synthesis error rate, these models generate the distributions of different kinds of errors, i.e., insertion, deletion, transition, and transversion (Figure 5). From Figure 5, it can be seen that, all linear models have positive slopes, which is quite obvious because the distribution of different kinds of errors increases with the increase of overall synthesis error rate. It can also be observed that, the linear model for deletion error has the maximum slope, while the linear model for the distribution of insertion error has the minimum slope. This is because, in order to develop these linear models, we used data from DNA synthesis technologies which had substantially high deletion rates (Figure 2a, 3, Figure 4a). On the other hand, insertion rates contributed the least among all types of errors in case of DNA synthesis technologies mentioned here. The distributions derived from these models were used to determine the type of error in step three.

**Figure 5.**
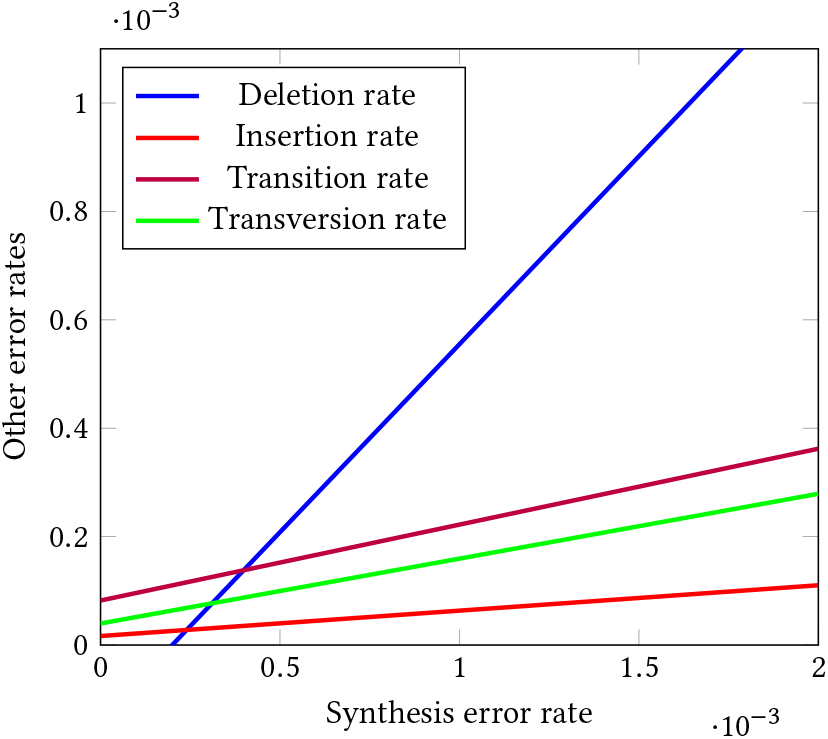
Linear models for determining distributions of different kinds of errors based on overall synthesis error rate.

Next, to determine the nucleotide that is likely to be involved in the selected type of error (step four), we needed to figure out the contributions of different nucleotide bases for different types of errors. Analogously, the distributions of the nucleotide bases in various kinds of errors were utilized from three DNA synthesis technologies mentioned above, in order to develop some linear models. Depending on the rate of a particular type of error, these models generate the distributions of four nucleotide bases for that error. These distributions were used in a stochastic model to select a particular nucleotide (step four). After that, a random bp location was selected within the DNA sequence, assuming that the error is likely to take place anywhere within the DNA sequence (step five). (Bayzid: this community may like passive voice sentence more than “we do” type sentences. So do this as much as possible. This is true for the entire paper.) Then, this error was directly introduced if the selected bp location contained the selected nucleotide (step six, seven). Otherwise, the neighbourhood of the chosen bp location was searched until one was found (step nine, ten).

(Bayzid: shouldn’t it be an option of custom error rate where we set error = 0? why do we need a separate option for error-free model?)Mahjabin: Sir, it is done Moreover, by setting the custom error rate to zero, users can simulate the ideal case, where no synthesis errors take place, i.e., the steps of synthesis module are bypassed altogether. This feature mainly allows us to evaluate the standalone performance of the other modules of DNA storage.

Finally, at the end of the steps in synthesis module, the error rates during this phase are written in a text file named *result*.*txt*. Also, the detailed information of each error (the nucleotide that participated in that error and error location) are recorded in a *log* file. Finally, the modified nucleotide sequence containing synthesis errors is passed on to next phase.

### 2.2 Preservation Module

One of the attractive features of DNA storage posits it as a longterm repository, able to retain data for over thousands of millions of years [19]. However, to make the retrieval of digital information possible from DNA storage even after millions of years, careful preservation strategies need to be considered because some finite degradation rate is always present in any sample regardless of storage conditions. Therefore, the integrity and fidelity of DNAs should be carefully maintained during storage period. In most instances, DNA samples are stored at very low temperatures or in liquid nitrogen. However, this would cause high maintenance cost for hundreds of millions of samples [6]. Therefore, for some time, micro-organic spores were being considered as suitable media for DNA storage for their ability to withstand hostile conditions that may arise during long term preservation periods [19, 57, 69]. However, spore resistance of many microorganisms remains unknown as well as poor spore revival most often results in incomplete retrieval of stored information [19]. After that, the next viable option was dehydrated DNAs stored at room temperatures for they greatly reduced cost and increased convenience [6]. The work of Bonnet *et al*. established that solid-state DNAs can be stored in room temperature if completely protected from water and oxygen [14]. However, it has been observed that DNA secondary structure undergoes denaturation and other changes upon dehydration or at lower levels of humidity [9, 30, 39]. Hydrolysis due to depurination, hydrolytic cytosine deamination, and oxidation contribute to the degradation of stored DNAs [28, 57, 92]. Among these factors, depurination is the hydrolytic cleavage of purine bases (adenine and guanine), which leads to both information loss and increased rates of chain break. This happens due to the absence of nucleotide bases and high rates of hydrolysis at apurinic sites [65]. It has been observed that dried state DNAs degrade spontaneously via depurination [71] though the rate is very low at room temperatures [14, 98]. A number of studies have established that the kinetics of DNA depurination reaction follows Arrhenius equation 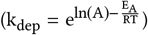 [8], i.e., the rate of DNA degradation rises exponentially with higher temperature [4, 14, 38, 59, 98]. In this equation, T is the absolute temperature measured in Kelvin, R is the molar gas constant (8.314 JK^−1^mol^−1^), E_A_ is the activation energy, i.e., the minimum energy required for DNA depurination, and A is the pre-exponential factor.

Besides, deamination [61] plays a major role in the dynamics of DNA degradation. In this process, cytosine spontaneously releases ammonia and changes into uracil via hydrolysis reaction. This results in miscoding lesions, i.e., differences in base-pairing in double-stranded DNAs (C →T and G→ A errors) [36, 53]. Though previous studies identified only age/ time as an important factor for DNA deamination reaction [83], Kistler *et al*. showed that cytosine deamination is strongly influenced by temperature as well as time. They proposed a simple linear thermal age model for cytosine deamination based on mammal bone samples available from published genomic data [53]. However, our analysis of this data revealed that temperature dependence of cytosine deamination fits exponential model (r^2^ = 0.122, P = 2.23 × 10^−6^) better than simple linear model (r^2^ = 0.068, P = 0.0005). Therefore, for our model, we used Arrhenius equation of temperature dependence 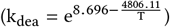 DNA deamination reaction. We found that cytosine deamination in mammal bone samples is associated with an activation energy of 39 kcal/ mol, which differs from the previously reported E_A_ in the work of Lindahl and Nyberg [60]. They estimated that the reaction is associated with an activation energy (E_A_) of 29 kcal/ mol. This difference might result from the fact that cytosine deamination in denatured DNA shows greater dependence at higher temperatures (70-95°C), but is less strongly dependent on temperature outside this range [60]. In their work, Lindahl and Nyberg explored the nature of cytosine deamination at higher temperatures (65-110°C), whereas, the reported deamination rates in mammal bone samples were within − 15 to 18°C.

Now, for the preservation module of our simulator, we tried to mimic DNA degradation in various natural and artificial storage media as well as its consequences using a number of mathematical models as described below.

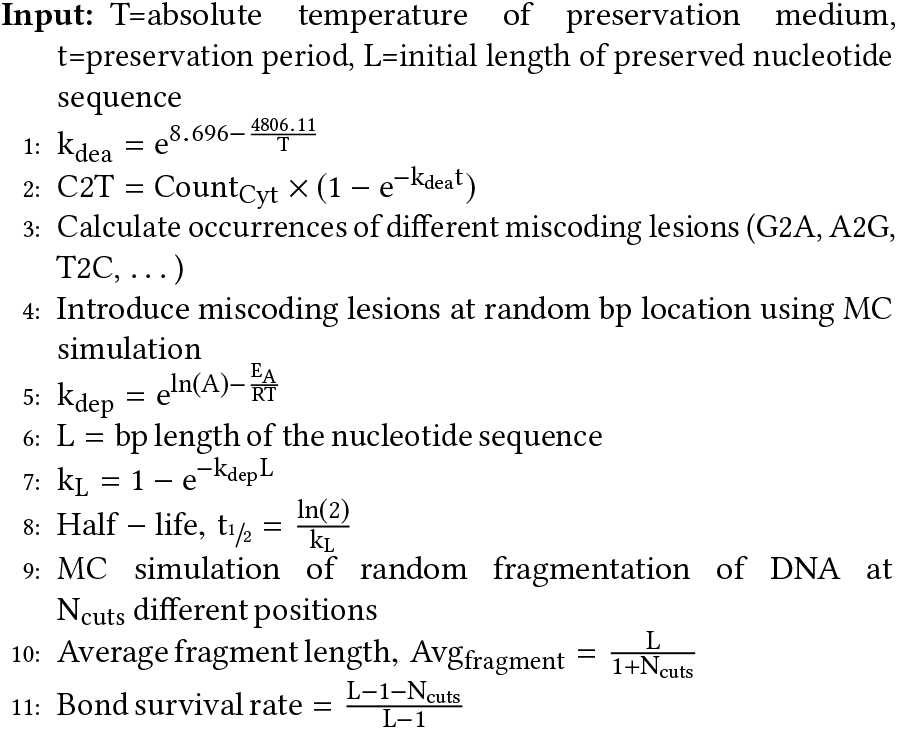

To begin with, the user needs to select a preservation medium and provide preservation period (t) as well as preservation temperature (T) to use this module. The user can provide temperature (T) either in Celsius or Kelvin. However, to use Arrhenius equation, we converted user provided temperature into Kelvin. On the other hand, the user may provide time (t) either in seconds, minutes, hours, days, weeks, months, or even years. Whatever be the unit of t, it is converted into seconds for further analyses.

First of all, we used Arrhenius equation to calculate cytosine deamination rate (k_dea_) at temperature T provided by the user (step one). For this purpose, we used our exponential model derived from the analysis of cytosine deamination rates in mammal bone samples at varying temperatures, used in the work of Kistler *et al*. [53]. Since all the reported deamination rates were in nt^−1^ year^−1^, the Arrhenius model of temperature dependence would generate k_dea_ in nt^−1^ year^−1^. So, we had to convert k_dea_ into nt^−1^ s^−1^ dividing it by 365× 24× 60 ×60 (the number of seconds in a year).

Next, we wanted to calculate the number of miscoding lesions caused by cytosine deamination using the formula: 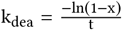,where t is time in seconds and x is damage occurrences per base sequenced [36, 53]. We can calculate x, i.e., damage occurrences per base sequenced by rearranging the equation as, 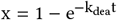.Now, to quantify the amount of miscoding lesions due to cytosine deamination for full-length nucleotide sequence, we multiplied per nucleotide damage occurrence, x with the number of cytosine (Count_Cyt_) sequenced from original DNA sequence. Hence, the amount of miscoding lesions due to cytosine deamination is, 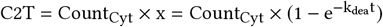 (step two).

Now, in case of cytosine deamination, if the DNA is sequenced from Light strand molecule, it will yield a C→ T miscoding lesion, whereas, the same event will appear as a G →A transition when sequenced from complimentary Heavy strand. Therefore, Hansen *et al*. suggested that miscoding lesions can be grouped into six complimentary and effectively indistinguishable pairs for difficulties associated with identifying the strand of origin of miscoding event [41]. These groups constitute of (A→ C, T→ G), (C→ G, G→ C), (A→T, T →A), (C →A, G →T), (A→ G, T→ C), and (C →T, G →A) miscoding pairs. We tried to quantify the occurrences of other types of miscoding lesions from the number of damage events caused by cytosine deamination (C2T). In this regard, we used empirical distribution of different types of damage events from the work of Gilbert *et al*. (Figure 6) [36]. In their work, they studied the nature of miscoding lesions, which constitute true damage events in ancient DNA. They demonstrated that C→ T and G→ A miscoding lesions constitute the overwhelming majority (88%) of miscoding lesions in ancient DNA. We used these distributions to calculate the occurrences of different miscoding lesions in the preserved nucleotide sequence over the course of preservation period, t (step three).

**Figure 6.**
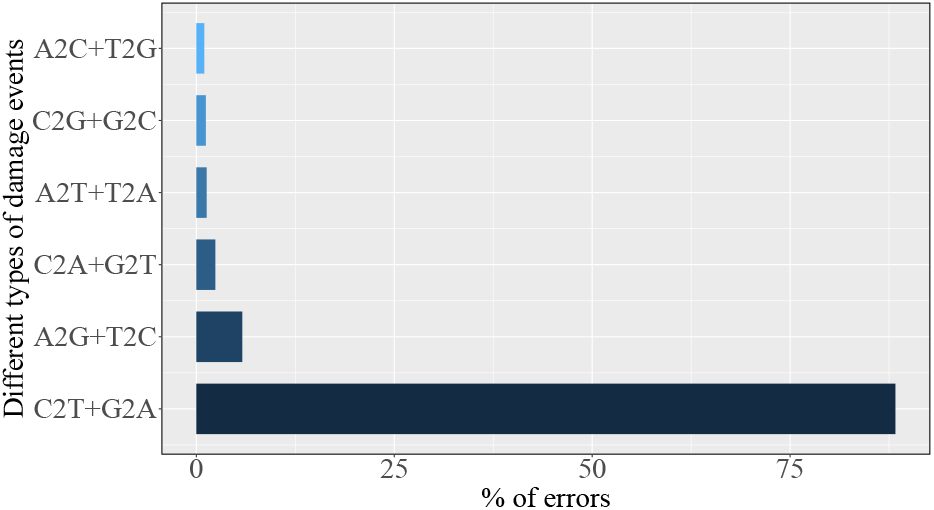
Empirical distribution of different types of damage events in ancient woolly Mammoth sequence from the work of Gilbert *et al*. [36].

Next, we ran Monte Carlo simulation to introduce all these damage events in the preserved nucleotide sequence. It has been noted that deamination errors do not affect the observed difference in DNA concentration [80]. Therefore, we can assume that the probability of damage events due to deamination is comparable across the length of random DNA sequences. For this, we generated random sites within preserved nucleotide sequence to introduce all the miscoding lesions one by one (step four).

After introducing errors caused by DNA deamination reaction, our preservation pipeline moves towards errors caused by DNA depurination reaction. In this regard, we used Arrhenius equation, (k_dep_ nt^−1^ s^−1)^to calculate DNA depurination rate in step five. For this purpose, we used empirical data of DNA depurination rates at varying temperatures in different types of storage media as mentioned below.

- **Solid-state DNA:** Bonnet *et al*. explored degradation of solidsate DNA at different temperatures to measure its stability as room temperature storage [14]. Since DNA degradation rate is too low to be conveniently measured at room temperature, they ran DNA chain-breaking kinetics at temperatures ranging from 70°C − 140°C. They reported DNA chain-breaking rate at 100°C as 10^−9^ nt^−1^ s^−1^ and extrapolated chain-breaking rate at room temperature (25°C) as 4.6×10^−15^ nt^−1^ s^−1^. Using these values in Arrhenius equation, we found that the activation energy (E_A_) of DNA depurination in solid-state DNA is 153792 J mol^−1^ and logarithm of pre-exponential factor (ln (A)) is 29.03 nt^−1^ s^−1^.
- **DNA on filter cards:** Burgoyne patented a solid medium for DNA storage utilising a solid matrix whose composition protects against degradation of DNA incorporated into the matrix [16]. This matrix comprises a solid support using absorbent cellulose-based filter paper, or micro-mesh of synthetic plastics material with DNA-protecting compound absorbed onto the support. Robert N. Grass and his team tested the stability of this dry storage technology at varying temperatures in their work [38]. They estimated that activation energy (E_A_) of DNA depurination on this filter card is 155632 J mol^−1^ and logarithm of pre-exponential factor, ln (A) is 43.44 s^−1^.
- **DNA in biopolymeric storage matrix:** Biopolymeric storage matrix is a cost-effective, environment friendly, room temperature DNA storage. It protects stored DNAs from heat and UV light by mimicking the anhydrous vitreous state of DNA in seeds and spores [99]. Grass *et al*. explored depurination of DNA in this storage matrix at various high temperatures [38]. They found that activation energy (E_A_) of DNA depurination reaction in biopolymeric storage matrix is 152706 J mol^−1^ and logarithm of pre-exponential factor (ln (A)) is 42.18 s^−1^.
- **DNA encapsulated in silica:** Grass *et al*. developed a protocol for encapsulating DNAs into amorphous silica spheres that mimics the protection of nucleic acids within ancient fossils [76]. In this approach, the nucleic acid molecules are hermetically sealed within glass spheres, which protect them from chemical attacks and thereby, enable them to withstand high temperatures and aggressive radical oxygen species. To measure its stability as room temperature storage, Grass *et al*. performed accelerated aging tests on DNAs encapsulated in silica [38]. From their experiments, they calculated activation energy (E_A_) of DNA depurination reaction in silica as 155221 J mol^−1^ and logarithm of pre-exponential factor (ln (A)) as 42.12 s^−1^.
- **Fossils:** DNA has the greatest chance of survival in ancient fossil bones when encapsulated within collagen structures and crystal aggregates, which protect DNAs from humidity and other environmental factors [31, 82]. Earlier findings from ancient moa bones revealed that DNA decay in ancient fossils follows exponential model [4]. Allentoft and his group analyzed mitochondrial DNAs from 158 radiocarbon-dated bones of extinct New Zealand moa and determined the activation energy (E_A_) of DNA decay in this natural storage as 127000 J mol^−1^. They also measured logarithm of pre-exponential factor ln(A) as 41.2 year^−1^.

Now, we can see that the reported unit of pre-exponential factor varies from experiment to experiment. Therefore, we wanted to take all of them into a single uniform unit, nt^−1^ s^−1^. Firstly, the activation energy and pre-exponential factors of DNA depurination reaction in filter cards, biopolymeric storage matrix, and synthetic silica spheres were measured for 158 bp long nucleotides

[38]. Therefore, the decay rate calculated using these values at step five would result in decay rate for 158 bp DNA, i.e., k_158bp_ measured in s^−1^. So, if the user selects one of these storage media, we need to convert this decay rate, k_158bp_ from s^−1^ to average per nucleotide decay rate, k_dep_ in nt^−1^ s^−1^. Now, it has been observed that DNA depurination reaction causes cleavage of strands which results in fragmented DNAs [58, 59, 75, 83]. This random fragmentation of DNA exhibits negative exponential correlation between fragment length and number of available copies [1, 23]. Therefore, Allentoft and his team reported the rate of full-length DNA degradation k_L_ in terms of Poisson distribution [4]. According to Poisson distribution, if average decay rate is k_dep_ nt^−1^ s^−1^, then probability of no decay per unit time across the length (L) of nucleotide sequence is 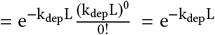. Hence, the probability that DNA depurination would occur across the length of nucleotide sequence or the rate of DNA depurination reaction for a given nucleotide sequence of length L is, 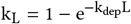.On the other hand, Robert N. Grass and his team showed that length-dependent DNA degradation kinetics follows linear model [65] assuming that the rate of full-length DNA degradation (k_L_) is the sum of decay rates of individual nucleotides, i.e., k_L_ = k_dep_L. Now, the length-dependent DNA degradation kinetic based on Poisson distribution also approximates to a linear model because the higher order terms tend to be negligible for the lower magnitude of decay rate k_dep_ at low preservation temperature.

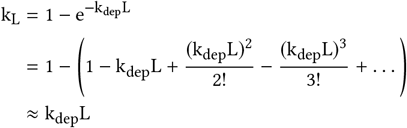

Hence, we used this exponential model 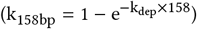 to get average per nucleotide decay rate (k_dep_) from k_158bp_ decay rates in filter cards, biopolymeric storage matrix, and synthetic silica fossils. On the other hand, the pre-exponential factor for DNA depurination in ancient moa fossils was reported in year^−1^. As a result, the calculated depurination rate k_dep_ would be in nt^−1^ year^−1^. So, we converted k_dep_ to nt^−1^ s^−1^ in case a user selects this natural preservation medium.

After calculating average per nucleotide depurination rate k_dep_ (nt^−1^ s^−1^), we converted it to depurination rate for full length nucleotide sequence, k_L_ (s^−1^) (step seven). Now, with full length DNA depurination rate at hand, we wanted to calculate half-life (t_1/ 2_) for preserved nucleotide sequence. It has been observed that DNA decay rate follows first-order kinetics [4, 38]. Therefore, at step eight, we used the equation of half-life for first-order chemical kinetics, 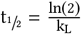

Next, we wanted to calculate how much the preserved nucleotide sequence would decay due to DNA depurination reaction over the course of preservation period, t. As we have already mentioned, DNA decay follows exponential relation, therefore, the number of nucleotides that would survive decay over the period t would be 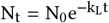 [4]. Here, N_0_ and N_t_ are the number of nucleotides at the beginning of the preservation period and after time, t respectively. In our case, N_0_ = L, the length of nucleotide sequence that need to be preserved, k_L_ is the decay rate for full length nucleotide sequence, measured in s^−1^, and t is the preservation period.

Now, the number of decayed molecules after period t would be: 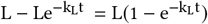. Since this number of purine bases would be released from the preserved nucleotide sequence, it would result in random fragmentation of DNA as mentioned earlier. Therefore, the expected number of cuts or breaks into preserved nucleotide sequence over the course of preservation period would be 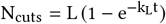 (step nine). After that, we ran Monte Carlo simulation to model this random fragmentation of DNA decay, assuming that the probability of breaking a strand at any nucleotide base is equivalent (step ten). As the probability of breaking a strand at any position is identical, the model generates a random internal position within the preserved nucleotide sequence where the strand breaks. A cut breaks the strand into two fragments, therefore, N_cuts_ would result in a total of 1 + N_cuts_ fragments. Hence, the average fragment length would be 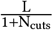 (step eleven). On the other hand, L nucleotide bases are linked via L− 1 phosphodiester bonds. Now, the expected number of bonds that would break due to depurination is N_cuts_. Therefore, bond survival rate for the preserved nucleotide sequence is 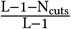.Finally, at the end of the preservation module, the random fragments generated by this process are passed to next module.

### 2.3 Sequencing Module

Last but not the least, comes the sequencing module that controls successful retrieval of stored information in DNA storage. There are many state of the art DNA sequencing technologies available around to this day, e.g., Sanger sequencing, next-generation sequencing, nanopore sequencing, etc. All of them try to determine the order of four nucleotide bases (A, T, C, G) in a given nucleotide sequence. In order to read a given piece of information in the form of DNA, first of all, we need to make multiple copies of it and break them into different fragments. These small fragments are then read and finally, assembled together to get the complete picture of the stored information. This whole pipeline of reading back DNA comes with lots of computational costs and associated complexities. However, with the advancement of new forms of technology, not only the pipeline of DNA sequencing has become faster and more efficient but also the cost associated with it has greatly reduced over the past decade. As of 2019, the cost of sequencing 1Mb of DNA has dropped to a mere 1.4 cents and the cost of sequencing a human-sized genome hovers around $1, 300 [48].

Besides various cost-effective technologies for DNA sequencing, in recent years, many computational tools have been developed for sequencing genome data. For example, MetaSim [79], Grinder [7], Mason [43] etc., simulate Sanger sequencing via empirical error models and other sophisticated design capabilities. Among nextgeneration sequencing simulators, there are MetaSim based on empirical error models of sequence reads [79], machine-learning based BEAR [51], DWGSIM - a whole genome simulator [44], EAGLE [73], FASTQSim [87], and so on. These computer-aided softwares facilitate the assessment and validation of various biological models and also help our understanding of specific data sets [26]. We may utilize computer simulation as a guide for developing new computational tools [46], proposing new hypotheses [42], designing sophisticated sequencing technologies [87, 88], and also for verifying the correctness of an assembly [54]. Now, for the sequencing module of our simulator, we tried to model reading errors based on empirical data as they represent real scenarios though the true process underlying them is usually unknown. The fundamental steps of our reading pipeline (sequencing module) are similar to that of writing process (synthesis module) and are as follows:

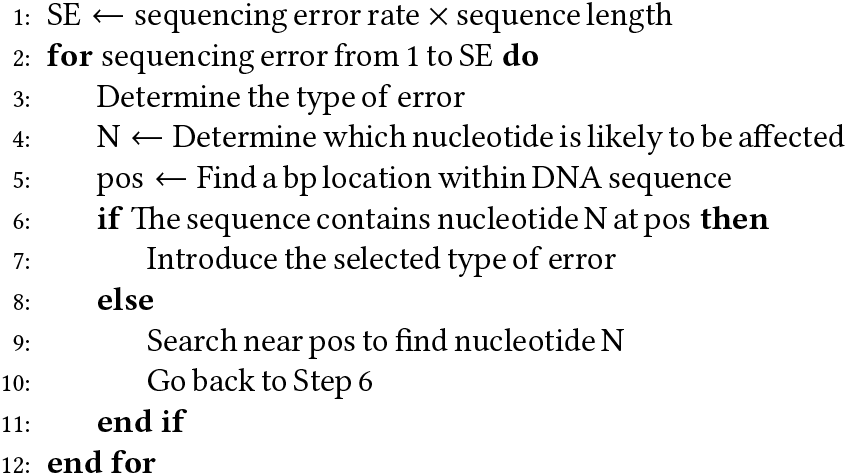

In the following we discuss four state of the art DNA sequencing technologies that we modeled in our simulator.

#### (1) Sanger Sequencing

Sanger sequencing is a “first generation” DNA sequencing method developed by two time Nobel Laureate Frederick Sanger and his colleagues in 1977 [70]. It is a chain termination method for determining the nucleotide sequence of a given piece of DNA. This technology has been able to achieve read lengths of up to 700− 1000 bp [90] and per bp accuracy as high as 99.99% [89].

Though Sanger sequencing produces limited data sets, it serves as an orthogonal method for confirming sequence variants identified by next generation sequencing (NGS) and also provides a means to patch the coverage of poorly-covered regions by NGS [40]. Also, it is suitable for measuring the fidelity of low-fidelity polymerases without any need of proofreading [77].

For our design of this part of sequencing module, we used empirical error distributions from the work of Potapov and Ong [77], where they.

## 3. EVALUATION

To measure how closely different modules of our simulator captured the nature, randomness, biases, and eccentricity of errors at different phases of DNA storage, we compared the output from each of these modules with different experimental outcomes from which we adopted empirical distribution of errors into different stochastic models. More importantly, we also compared our outcomes with different *in vitro* experimental results, which we did not use to model the error distributions in SEEDS, to see how our stochastic models compare with new and unseen experiments. We were not only interested to find out the degree to which our models captured the chaotic nature of errors in DNA but also to figure out how closely our models reflected the metabolic nature of DNA during preservation. The performance of different modules of our simulator are presented in detail as follows.

### 3.1 Performance of synthesis module

First, we compared the output of synthesis module with error distributions of different synthesis technologies from which we integrated error rates in our simulator. In this regard, we used DNA sequences of different gene constructs from the work of Quan *et al*. [78] as input nucleotide sequence of our simulator. We did not incorporate any data from their work in our stochastic error models. To evaluate the sole performance of synthesis module, we passed the input nucleotide sequence only through this module of our simulator while setting “no error” option for both preservation module and sequencing module. We compared the average outcome over 100 iterations against the error distributions of different state of the art DNA synthesis technologies.

First of all, we compared the outcome of ‘*de novo* DNA synthesis’ module of our simulator with experimental outcomes from the work of Carr *et al*. [17] from which we incorporated empirical error distributions into this part of synthesis module. From Figure 7a, we can see that the simulated error rates varied little from that of experimental error rates. The overall synthesis error rate of our simulator differed from that of wet-lab implementation of Carr *et al*. [17] by a degree of 1 in 10000. This is because we used normal distribution to generate random error rates to account for varying synthesis error rates from experiment to experiment using this technology. Accordingly, the error rates for different types of errors (deletion, insertion, transition, and transversion) also varied because they depended on the overall error rate. However, none of these error rates varied more than a few hundred thousandths.

**Figure 7.**
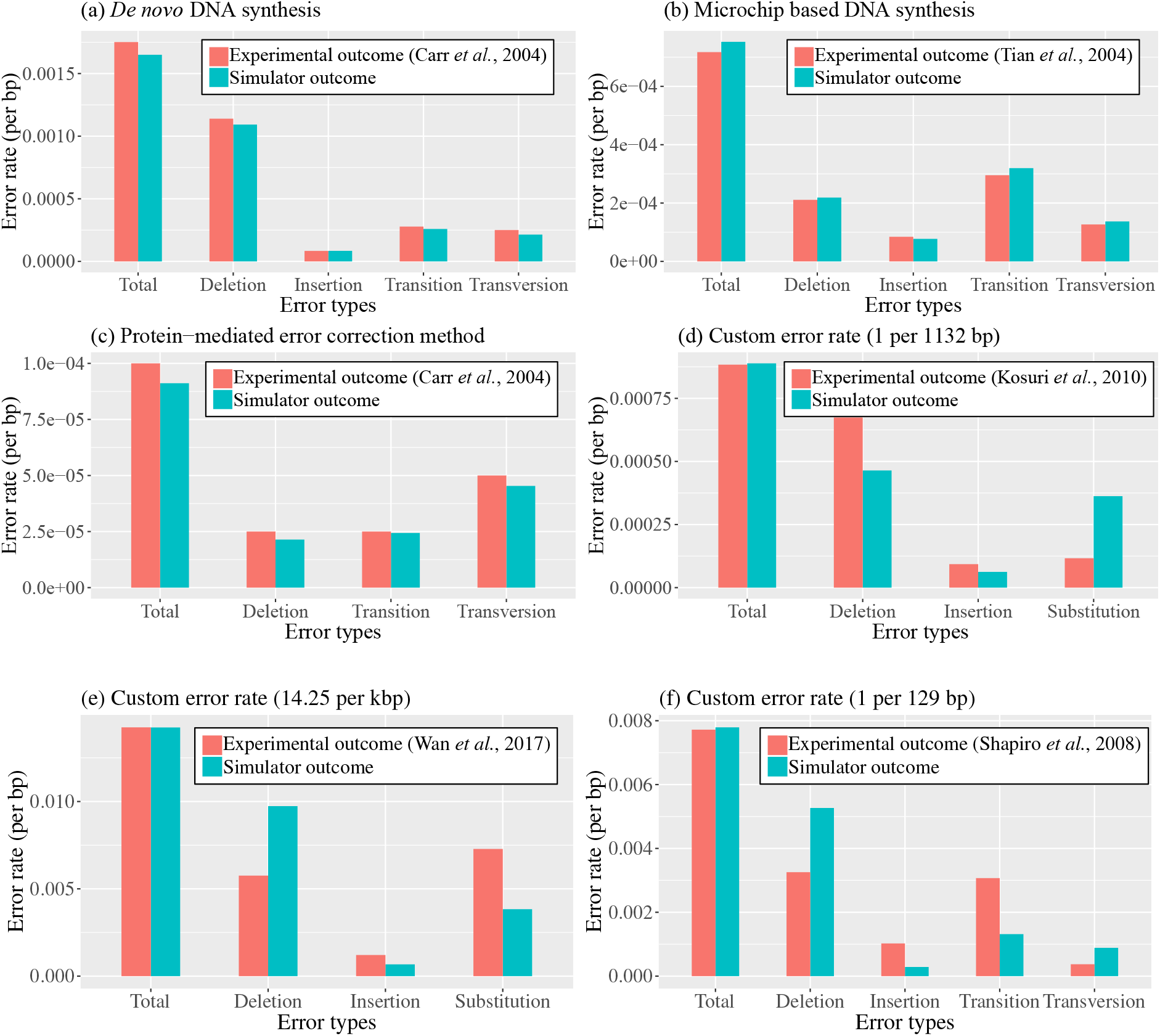
Evaluation of outcomes of synthesis module with different experimental outcomes. Here, simulated error rates are compared against error distributions from different DNA synthesis technologies.

Secondly comes the ‘microchip based DNA synthesis module’ of our simulator. We compared its outcome with wet-lab implementation of multiplex gene synthesis using programmable DNA microchips by Tian *et al*. [94]. From their work, we integrated the information about the distribution of different kinds of errors in microchip based DNA synthesis into our stochastic models. When we compared the error rates in our outcome with that of experimental outcomes, we found that the simulated error rates closely followed experimental error rates (Figure 7a). The overall error rate in our simulator varied from that of experimental outcome by a few hundred thousandths. This is because, we used random synthesis error rates generated by normal distribution of varying error rates in oligonucleotides synthesized on microchips in various experiments. Therefore, our overall synthesis error rate varied from the reported error rate (1 in 1394 bp) in the work of Tian *et al*. [94]. Analogously, the rates of other kinds of errors also varied for their dependence on overall error rate. However, their difference with experimental error rates were in the range of a few millionths to a few hundred thousandths.

The third synthesis technology from which we collected information for developing our stochastic models was ‘protein-mediated error correction for *de novo* DNA synthesis’. For this part of synthesis module, we compared simulator outcomes with reported error rates in the work of Carr *et al*. [17] from which we incorporated error rates into our model. From Figure 7a, we can see that the simulated error rates followed the experimental error rates in close steps. The difference between experimental error rate and average simulated error rate over 100 iterations was only a few millionths per bp because we used overall synthesis error rate from experimental data directly in our model. Even, the difference between other types of errors were in the range of one in ten-millionths to a few millionths.

Apart from verifying the performance of our simulator against the experimental settings from which we adopted our stochastic error models, our main challenge was to incorporate robustness into our models to handle arbitrary synthesis error rates. Therefore, we checked the performance of ‘custom error rate’ feature of our synthesis module against relevant experimental results from the works of Kosuri *et al*. [55], Wan *et al*. [100], and Shapiro *et al*. [106].

Sriram Kosuri, George Church, and their team used high-fidelity DNA microchips known as OLS (Oligo Library Synthesis) pools to achieve a better starting point for more scalable microchip-based DNA synthesis. They designed two OLS pools with oligonucleotides of varying lengths and one of them used GFP35 as independent assembly subpool that encoded GFPmut3b. For this GFP35 synthesis reaction, they found an error rate of 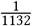 per bp after sequencing on Illumina. Since next generation sequencing platforms, such as Illumina are more error prone than Sanger-based sequencing approach, they used only high-quality reads (> 99.9% accuracy) for mapping. Therefore, we used their reported error rate as custom error rate 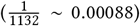 in our synthesis module and set ‘no error’ for both preservation and sequencing modules because they did not have any preservation phase for the synthesized oligonucleotides and used high-quality sequencing reads for mapping with original sequence. With these settings, we again passed our input nucleotide sequences through different modules for over 100 iterations. The results revealed that the overall synthesis error rate from our synthesis module closely matched with the original reported error rate (Figure7a) because we used this rate directly to calculate the number of possible synthesis errors in step one. However, the rates of different kinds of errors varied from the reported error rates in original GFP35 synthesis. For example, deletion rates were higher during GFP35 synthesis, whereas, our linear models introduced lower deletion rates by an amount of 2 errors less per 10000 bp. On the other hand, the simulated insertion error rate was pretty close to that of the actual error rate (difference 0.00003 errors ber bp). And finally, our linear models introduced higher (2 errors more per 10000 bp) substitution rates (transitions and transversions), whereas, these errors were actually lower in the original experiment.

Next, we evaluated the performance of custom error module of our simulator with the work of Wan *et al*. [100]. They used highly error-prone microchip-synthesized oligo pools for *de novo* synthesis of genes associated with a biological pathway. Using a combination of error removal techniques they were able to reduce error rates from 14.25 /kbp to 0.53/kbp. We used their reported error rate of 14.25/kbp without any error correction mechanism for evaluating the performance of our synthesis module. We again found that the overall synthesis error rate was identical with that of the reported error rate (Figure 7b). Analogously, the error rates for other types of errors varied from that of the reported error rates. For instance, in absence of error removal, deletion errors were abundant (0.0058 errors per bp) within synthesized fragments. However, the deletion rates introduced by our models were higher (0.0097 errors per bp, i.e., 39 deletions more per 10000 bp) than that of the actual deletion error rate. On the contrary, the simulated error rates for insertion closely matched with that of the original error rate (difference ∼0.0005 errors per bp). And in case of substitutions, our models introduced lower errors (35 substitutions less per 10000 bp) than that of the actual amount of substitutions.

Lastly, we used error distributions from the work of Shapiro *et al*. [106] to evaluate the robustness of our synthesis module. They used single molecule PCR-based technique as an alternative for in vivo cloning of DNA, which is one of the bottlenecks of DNA writing (synthesis). Integrating this rapid, high-fidelity in vitro procedure into DNA synthesis and error correction mechanisms, they were able to efficiently construct in vitro DNAs from unpurified oligos. Without any error correction mechanism, they reported an error rate of 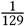 per bp for in vitro clones of GFP constructs. We used this error rate 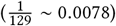 as custom error rate in synthesis module of our simulator and evaluated the outcomes with experimental results. We found that the overall error rates matched closely (Figure 7b) but the rates for other types of errors differed from that of the experimental errors. For example, the in vitro clones of DNAs in their experiment had a deletion rate of 0.0032 per bp, whereas, our linear models introduced an error rate of 0.0053 per bp, which means in our simulator 21 bases will be deleted more per 10000 bp. In case of insertion, our simulated error rates trailed behind the reported error rates with a narrow margin of only 7 insertions per 10000 bp. On the other hand, the synthesized clones had higher transition rate than that predicted by our stochastic error model (18 errors less per 10000 bp). Finally, the amount of transversion errors introduced by our linear model was quite close to that of the actual errors, only 5 transversions more per 10000 bp.

So, we can see that our naive linear models, for the distributions of different kinds of errors, fared quite well in the face of arbitrary synthesis errors from different experimental settings from which we used no data to design our stochastic models.

### 3.2 Performance of preservation module

We evaluated the outcomes of our preservation module based on two criteria: (i) random fragmentation of DNA caused by DNA depurination reaction and (ii) characterization of miscoding lesions due to DNA deamination reaction. Following is a brief description of the findings available from our preservation module.

#### ▪ Temperature vs DNA Half-life

As it has been already mentioned, DNA decay via depurination follows first-order chemical kinetics and therefore, DNA has a half-life. However, depurination of DNA is affected by the nature of preservation medium. Therefore, to visualize the stability as well as fidelity of preserved nucleotide sequence in various storage media, we plotted DNA half-life against temperature at varying storage conditions (Figure 8). In this regard, we took a random sequence of length 158 bp and measured its half-life over a wide range of temperatures (from − 20° to 80° C) by simulating depurination under various storage conditions. We selected this particular length in order to compare the outcome of our model with that of original half-life of 158 bp DNA in the wet-lab implementation of Grass *et al*. [38]. We carried out 100 simulations of DNA depurination reaction for a particular medium and a given storage temperature.

**Figure 8.**
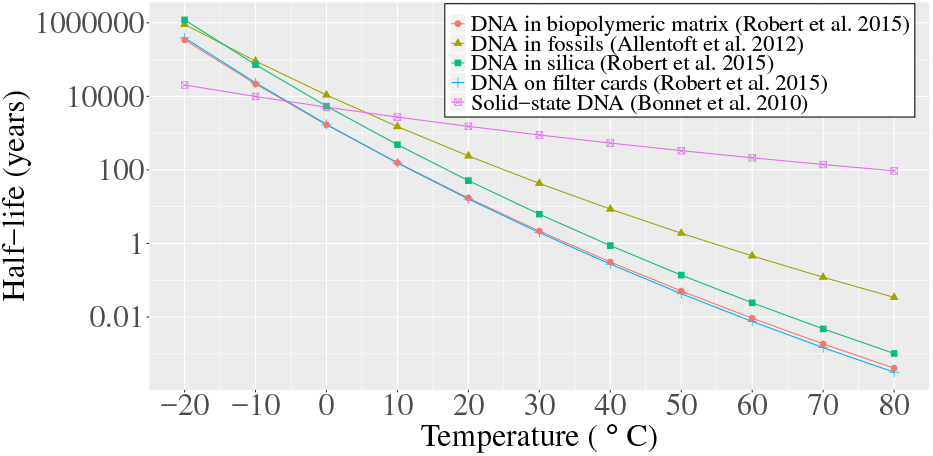
Half-life of 158 bp DNA stored in different storage media across a wide range of temperatures. The half-lives have been calculated following first-order chemical kinetics of DNA depurination reaction in different storage media as reported in previous literature. Here, the half-life is reported in log-scale.

It can be observed that, in terms of longevity, synthesized DNAs can be best preserved in solid-state in the absence of water and oxygen. However, previous studies have shown that upon dehydration, natural DNA undergoes reversible denaturation at room temperature [14]. Therefore, the next viable option could be ancient fossil bones. This is the only natural storage medium reported in our preservation module. The apatite/ collagen structures [31] and crystal aggregates [82] present within bones protect the solid DNAs from humidity and other environmental factors. Similar performance might be achieved via encapsulating DNA within inorganic silica particles as observed from the wet lab implementation of Grass *et al*. [38]. However, it is worth noticing from the outcomes of our simulator that though the stability of DNA within silica particles lags behind than that of the DNAs in ancient fossil bones at higher temperatures, their performance coincide at lower temperatures, i.e., at least as low as − 10° C. This indicates that by storing synthesized DNAs in inorganic silica particles at sufficiently lower temperatures, it might be possible to preserve them for thousands to millions of years.

Moreover, the other two storage media included in our module, i.e., DNA in biopolymeric matrix [99] and DNA on filter cards [16] perform similarly in terms of longevity of DNA across a wide range of temperatures. Both of these storage media comprise matrices of various kinds. For instance, filter cards consist of a synthetic polystyrene or cellulose-based solid matrix that provides support for the stored DNA [16], whereas, Biomatrica’s DNA SampleMatrix is based on glass polymers that shrinks and protects stored DNAs from various harmful external factors [99]. However, the stability of stored DNAs within both of these modules lags behind that of DNAs encapsulated in inorganic silica particles. This is because, among all the artificial storage media reported in our preservation module, silica-based inorganic storage medium has lowest level of local water concentration as well as exceptional stability against oxidation [38]. These attributes make it potentially more suitable compared to other storage media for long-term preservation of DNA.

#### ▪ Fragmentation of DNA

The role of depurination in DNA fragmentation is well-documented in previous research studies [58, 59, 75, 83]. DNA fragmentation theory proposes that the number of available copies decreases exponentially with fragment length [1, 4, 23, 84]. Therefore, we simulated (100 times) how a random sequence of 1000 bp would fragment over 100,000 years at 10°C preservation temperature across different media. The outcome of our simulation (Figure 9) confirms the exponential relationship between copy number and fragment length as posited by DNA fragmentation theory. It is worth noticing that the proportion of fragments of varying lengths are similar across different storage media after the same preservation period.

**Figure 9.**
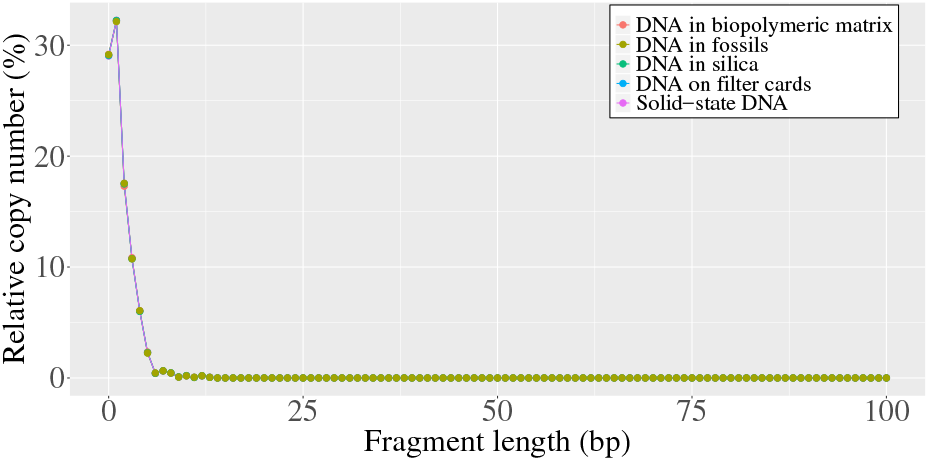
Exponential relationship between copy number and fragment length due to random fragmentation of DNA.

However, at the same preservation temperature (10°C) phospho-diester bonds remain intact longer when preserved in solid-state DNA compared to other storage media (Figure 10a). This is also evident from the average fragment length over time (Figure 10b).

**Figure 10.**
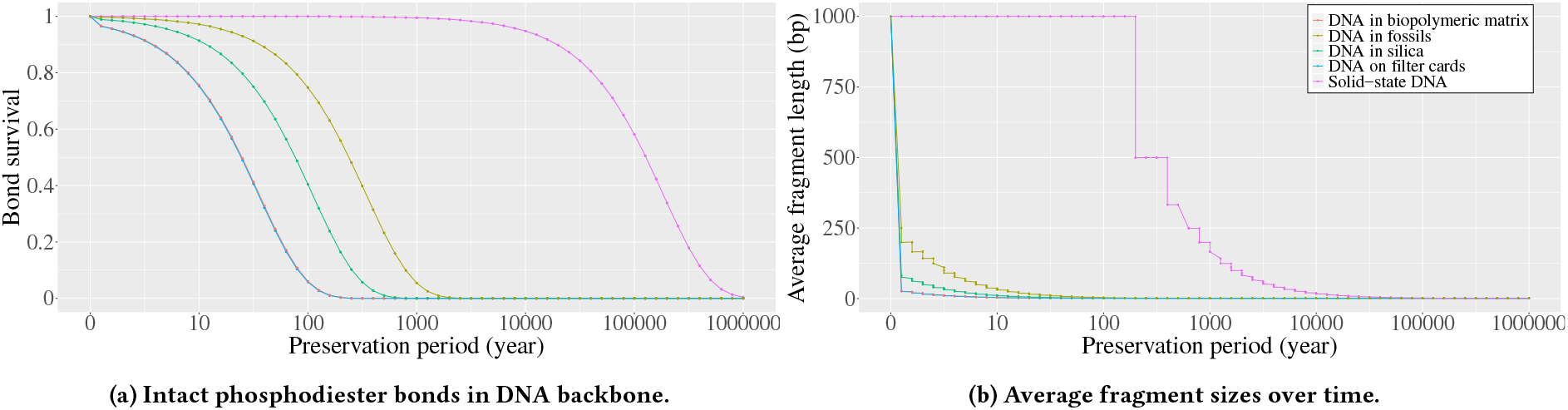
Bond survival and average fragment length for a 1000 bp DNA stored at different storage media at 10°C.

## 4 DISCUSSION AND CONCLUSIONS

In this paper, we proposed SEEDS, the first simulator that aims at mimicking the generation and propagation of different types of errors at various phases in DNA storage. SEEDS generates simulated error models by emulating the DNA storage process with technology- and preservation media-specific error models and decay rates. SEEDS uses various well know in vivo studies on DNA storage to generate appropriate statistical models to fit empirically derived errors. It covers various aspects of the entire pipeline in DNA storage including synthesis platform, preservation media, sequencing technologies, etc.

**Not sure if the next para is more suitable for introduction**…

The last few years have seen the emergence of a host of DNA storage technologies. However, in order to make it a reliable, cheap and practical means for digital data storage, various advanced techniques and error protection schemes need to be developed. It takes substantial amount of money and time to evaluate and compare various technologies. Therefor, it poses a barrier in assessing the performance of DNA storage techniques and thus the advancement in various algorithmic techniques and error correcting codes is riddled with non-trivial dependency problems on time and money. So it is imperative to have a reliable simulator that can mimic the error models in DNA storage.

SEEDS is the first of its kind and offers flexible settings to emulate various types of errors. SEEDS produces simulated errors that fit well with the data obtained in previous in vitro studies. <include discussion on how our simulator produced error models that fit well with unseen wet-lab data. By unseen, I mean those that were not included in modeling the error rates in SEEDS.> However, it can still be improved and extended in various directions. <List the limitations of SEEDs>.

The timing of this simulator seems appropriate as DNA storage techniques are receiving significant attention from the scientific community, various advanced techniques and error correcting codes are being developed which leads many to speculate on its near-term potential as a practical storage media, and finally evaluating the performance of various techniques and algorithms related to DNA storage is mostly dependent on extremely expensive and time consuming in vitro experiments. This study establishes a foundation for advancing our understanding of the impact of various factors on the error propagation in different DNA storage technologies. We believe SEEDS will be a powerful and useful tool for providing an in silico means for mimicking the error model in DNA storage under a wide range of practical and challenging model conditions. SEEDS will continue to evolve with the availability of new experimental data, and in response to new scientific findings – laying a firm foundation for DNA storage error simulators.

## Notes

### Competing Interest Statement

1. model modification and DL appliance

